# Inhibition of type III secretion system induced leukotriene B_4_ production by *Yersinia pestis*: A mechanism for early immune evasion

**DOI:** 10.1101/2023.03.13.532349

**Authors:** Amanda Brady, Amanda R. Pulsifer, Sarah L. Price, Katelyn R. Sheneman, Krishna Rao Maddipati, Sobha R. Bodduluri, Jianmin Pan, Shesh N. Rai, Bodduluri Haribabu, Silvia M. Uriarte, Matthew B. Lawrenz

**Author notes:** College of Medicine, University of Cincinnati, Cincinnati, United States (Jianmin Pan, Shesh N. Rai).

## Abstract

Subverting the host immune response to inhibit inflammation is a key virulence factor of *Yersinia pestis*. The inflammatory cascade is tightly controlled via the sequential action of lipid and protein mediators of inflammation. Because delayed inflammation is essential for *Y. pestis* to cause lethal infection, defining the mechanisms used by *Y. pestis* to manipulate the inflammatory cascade is necessary to understand this pathogen’s virulence. While previous studies have established that *Y. pestis* actively inhibits the expression of host proteins that mediate inflammation, there is currently a gap in our understanding of inflammatory lipid mediator response during plague. Here we use in vivo lipidomics to define the synthesis of lipid mediators of inflammation within the lungs during pneumonic plague. Interestingly, while we observed an early cyclooxygenase response during pneumonic plague, there was a significant delay in the synthesis of leukotriene B4 (LTB_4_), a pro-inflammatory lipid chemoattractant and activator of immune cells. Furthermore, in vitro studies with primary leukocytes from mice and humans further revealed that *Y. pestis* actively inhibited the synthesis of LTB_4_. Finally, using *Y. pestis* mutants in the Ysc type 3 secretion system (T3SS) and *Yersinia* outer protein (Yop) effectors, we demonstrate that leukocytes recognize the T3SS to initiate the synthesis of LTB_4_ rapidly. However, the Yop effectors secreted through the same system effectively inhibit this host response. Together, these data demonstrate that *Y. pestis* actively inhibits the synthesis of LTB_4_, an inflammatory lipid, required for rapid recruitment of leukocytes to the site of infection.

**Author Summary:** *Yersinia pestis*, the bacteria that causes plague, targets the host’s innate immune response to inhibit inflammation. Because the generation of this non-inflammatory environment is required for infection, we are interested in mechanisms used by *Y. pestis* to block inflammation. Lipid mediators are potent signaling molecules that regulate multiple host immune responses, including inflammation. While there have been studies on how *Y. pestis* blocks the proteins that mediate inflammation, there is a gap in our understanding of the inflammatory lipid mediator response during plague. Here we show that *Y. pestis* inhibits the production of one of these critical lipid mediators, leukotriene B4, by host immune cells. Furthermore, we identify both the signals that induce LTB_4_ production by leukocytes and the mechanisms used by *Y. pestis* to inhibit this process. Together, these data represent the first comprehensive analysis of inflammatory lipids produced during plague and improve our current understanding of how *Y. pestis* manipulates the host immune response to generate a permissive non-inflammatory environment required for bacterial colonization.

## Introduction

*Yersinia pestis*, a gram-negative facultative intracellular bacterium, is the causative agent of the human disease known as plague. Although typically characterized as a disease of our past, in the aftermath of the 3^rd^ plague pandemic, *Y. pestis* became endemic in rodent populations in several countries worldwide, increasing the potential for spillover into human populations through contact with infected animals and fleas (1-3). Human plague manifests in three forms: bubonic, septicemic, or pneumonic plague. Bubonic plague resulting from flea transmission arises when bacteria colonize and replicate within lymph nodes. Septicemic plague results when *Y. pestis* gains access to the bloodstream, either directly from a flea bite or via dissemination from an infected lymph node, and results in uncontrolled bacterial replication and sepsis. Finally, secondary pneumonic plague, wherein *Y. pestis* disseminates to the lungs via the blood, results in pneumonia that can promote direct person-to-person transmission via aerosols. While treatable with antibiotics, if left untreated, all forms of plague are associated with high mortality rates, and the probability of successful treatment decreases the longer initiation of treatment is delayed post-exposure (3-6). Regardless of the route of infection, one of the key virulence determinants for *Y. pestis* to colonize the host is the Ysc type 3 secretion system (T3SS) encoded on the pCD1 plasmid (5, 7). This secretion system allows direct translocation of bacterial effector proteins, called Yops, into host cells (5, 8, 9). The Yop effectors target specific host factors to disrupt normal host cell signaling pathways and functions (10-15). Because the T3SS and Yops are required for mammalian but not flea infection, the expression of the genes encoding these virulence factors are differentially expressed within these two hosts (5, 8, 16, 17). The primary signal leading to T3SS and Yop expression is a shift in temperature from that of the flea vector (<28°C) to that of the mammalian host (>30°C). During mammalian infection, *Y. pestis* primarily targets neutrophils and macrophages for T3SS-mediated injection of the Yop effectors (18-20). The outcomes of Yop injection into these cells include inhibition of phagocytosis as well as inflammatory cytokine and chemokine release required to recruit circulating neutrophils to infection sites (21-24). Importantly, previous work suggests that inhibition of neutrophil influx and establishing a non-inflammatory environment is crucial for *Y. pestis* virulence (25, 26). Therefore, defining the molecular mechanisms used by *Y. pestis* to subvert the host immune response is fundamental to understanding the pathogenesis of this organism. Moreover, defining the host mechanisms targeted by *Y. pestis* to inhibit inflammation can also provide novel insights into how the host responds to bacterial pathogens to control infection.

A cascade of events tightly regulates inflammation to ensure rapid responses to control infection and effective resolution after clearance of pathogens to limit tissue damage (27, 28). This inflammatory cascade is initiated by synthesizing potent lipid mediators and is sustained and amplified by the subsequent production of protein mediators (29, 30). Polyunsaturated fatty acid (PUFAs) derived lipid mediators are a family of lipids that critically enhance innate and adaptive immune inflammatory responses (28, 31). Of these, the eicosanoids, including the leukotrienes and the prostaglandins, are widely recognized for their role in influencing the inflammatory cascade during infection (29, 30). Leukotriene B4 (LTB_4_) is rapidly synthesized from arachidonic acid upon activation of 5-lipoxygenase (5-LOX), 5-LOX activating protein (FLAP), and LTA_4_ hydrolase (32). Previous work has established that LTB_4_ is essential for rapidly initiating the inflammatory cascade via engagement with the high affinity BLT1 receptor on resident effector leukocytes (29, 30, 33-36). BLT1 engagement promotes chemotaxis and stimulates effector cells to express and release pro-inflammatory cytokines that lead to chemokine production (37). These chemokines promote the recruitment of circulating leukocytes to the infected tissue (30). Importantly, because of its critical role in initiating the inflammatory cascade, disruption in the timely production of LTB_4_ can slow the subsequent downstream release of cytokines and chemokines and the ability of the host to mount a rapid inflammatory response required to control infection.

Despite active proliferation of *Y. pestis* within the lungs in the mouse model, previous studies have reported an absence of pro-inflammatory cytokines, chemokines, and neutrophil influx for the first 36 hours of primary pneumonic plague (11-15). This phenotype dramatically differs from pulmonary infection with attenuated mutants of *Y. pestis* lacking the T3SS or Yop effector proteins or by other pulmonary pathogens, such as *Klebsiella pneumoniae*, which induce significant inflammation within 24 hours of bacterial exposure (11-15). Surprisingly, despite the importance of lipid mediators in initiating and defining the inflammatory cascade in response to infection, the role of inflammatory lipids during plague has yet to be defined. In this study, we provide the first lipidomic profile of host inflammatory lipids during the initial 48 h of pneumonic plague and demonstrate a dysregulation in the production of LTB_4_ by *Y. pestis*. We further show that while leukocytes can quickly initiate the synthesis of LTB_4_ in response to the *Y. pestis* T3SS, the bacterium effectively inhibits the synthesis of this critical lipid mediator via the action of multiple Yop effectors secreted via the same T3SS. Together these data demonstrate active inhibition of LTB_4_ production by *Y. pestis*, providing new insights into the interactions between *Y. pestis* and host innate immune cells. Further, these data suggest that modulation in the production of host inflammatory lipids is an additional virulence mechanism used by *Y. pestis* to inhibit the rapid recruitment of immune cells needed to control infection.

## Results

### LTB_4_ synthesis is blunted during pneumonic plague

Despite the critical role lipid mediators play in the induction of inflammation within the host, the synthesis profile of lipid mediators during pneumonic plague has yet to be defined. Therefore, to establish the kinetics of lipid mediator production during pneumonic plague, C57BL/6J mice were intranasally infected with fully virulent *Y. pestis*, and lungs were collected at 6, 12, 24, 36, and 48 h post-infection. Total lipids were extracted from homogenized tissues with methanol, and 143 host inflammatory lipids were quantified by LC-MS and compared to naïve lungs (38). We observed significant changes in the synthesis of 63 lipids during infection, including lipids generally considered to be pro-inflammatory (18 lipids), anti-inflammatory (41 lipids), or pro-resolving (4 lipids) (S1 Table, S1 Fig., S3 Fig). However, it is important to note that categorizing inflammatory lipids is not simple, and many can have both pro- and anti-inflammatory properties depending on the lipid concentrations and the cell types that interact with the lipid (39, 40). Interestingly, we observed significant differences in the synthesis of the eicosanoids produced via the cyclooxygenase and lipoxygenase pathways (Fig. 1). While the synthesis of several of the cyclooxygenase-dependent prostaglandins increased by 6 h post-infection and remained elevated for 24-48 h (Fig. 1A-E), synthesis of LTB_4_ was not significantly elevated until 48 h post-infection (Fig. 1F). This directly correlated with the absence of significant amounts of the LTB_4_ degraded byproduct 20-hydroxy LTB_4_ at the same timepoints (Fig. 1G). However, we observed a significant increase in 5-HETE as early as 6 h post-infection (Fig. 1H), which can result if 5-LOX does not complete the synthesis of arachidonic acid to LTA_4_ (LTA_4_ is the precursor of LTB_4_) (41, 42). Interestingly, we did not observe significant synthesis of 5-oxo-ETE (Fig. 1I), which is derived from 5-HETE, indicating that oxidation of 5-HETE by 5-hydroxyeicosanoid dehydrogenase (5-HEDH) was not occurring within the infected tissues. Together these data suggest that while the cyclooxygenase pathway is induced rapidly during *Y. pestis* infection, LTB_4_ synthesis is specifically blunted during pneumonic plague. As LTB_4_ is a potent mediator in inflammation (33), we sought further to investigate LTB_4_ in the context of *Y. pestis* infection.

**Fig 1.**
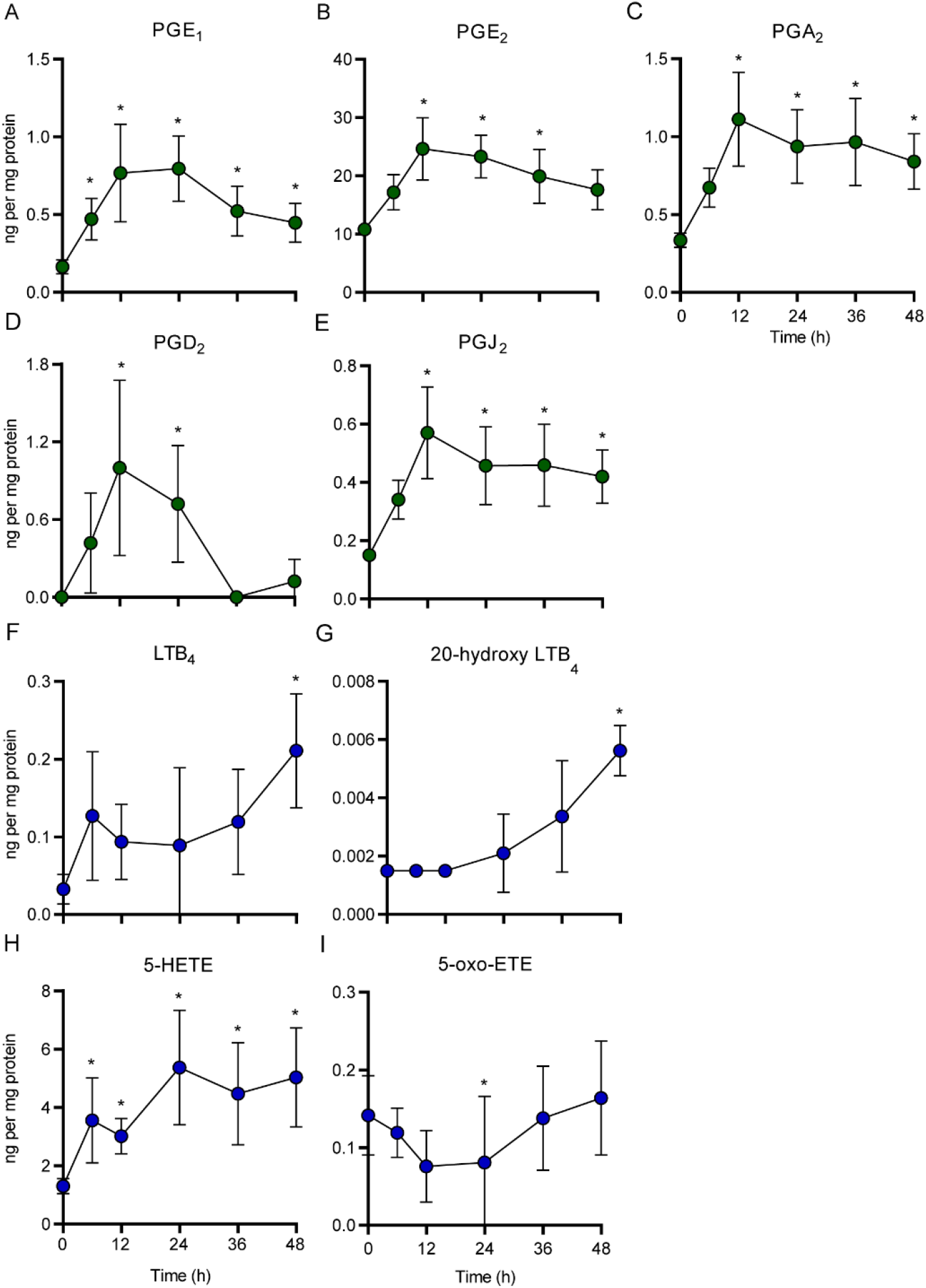
LTB_4_ synthesis is blunted during pneumonic plague. C57Bl/6J mice were infected with 10 x the LD_50_ of *Y. pestis* KIM5 and lungs were harvested at 6, 12, 24, 36, and 48 h post-infection (n=5). Lipids were isolated from homogenized tissues, quantified by LC-MS, and compared to concentrations from uninfected lungs (T=0). Green symbols = cyclooxygenase pathway; Blue symbols = lipoxygenase pathway. Median ± the range were compared by LIMMA-Moderated t-test; *=p≤0.001.

### BLT1^-/-^ mice are not more susceptible to pneumonic plague than C57BL/6J mice

LTB_4_ is recognized by the high-affinity G-protein coupled receptor BLT1, which is expressed primarily by innate and adaptive immune cells (36, 43). LTB_4_-BLT1 engagement leads to host inflammatory immune responses such as cytokine release, chemotaxis, phagocytosis, and reactive oxygen species (ROS) production that contribute to the clearance of pathogens (33, 44). Mice deficient in the expression of BLT1 cannot effectively respond to LTB_4_ signaling and are more susceptible to infections by bacteria and fungi (37, 45, 46). Because we did not observe LTB_4_ synthesis during the early stages of pneumonic plague, we hypothesized that BLT1^-/-^ mice would not be more susceptible to *Y. pestis* infection. To test this hypothesis, we intranasally infected BLT1^-/-^ mice with a fully virulent *Y. pestis* strain that carries a luciferase bioreporter that allows us to monitor bacterial proliferation and dissemination, in real-time, in live animals via optical imaging (47). Over the first 60 h of infection, we observed no difference in bioluminescent signal in the lungs of BLT1^-/-^ mice compared to wild type C57Bl/6J mice, indicating that the bacteria did not replicate faster in BLT1^-/-^ mice (Fig. 2A). BLT1^-/-^ mice also did not succumb to infection any quicker than the C57Bl/6J controls (Fig. 2B). These data demonstrate that the loss of LTB_4_-BLT1 signaling in BLT1^-/-^ mice has no impact on the infectivity of *Y. pestis*, and further supports our lipidomics data that LTB_4_ synthesis and signaling is disrupted during pneumonic plague.

**Fig 2.**
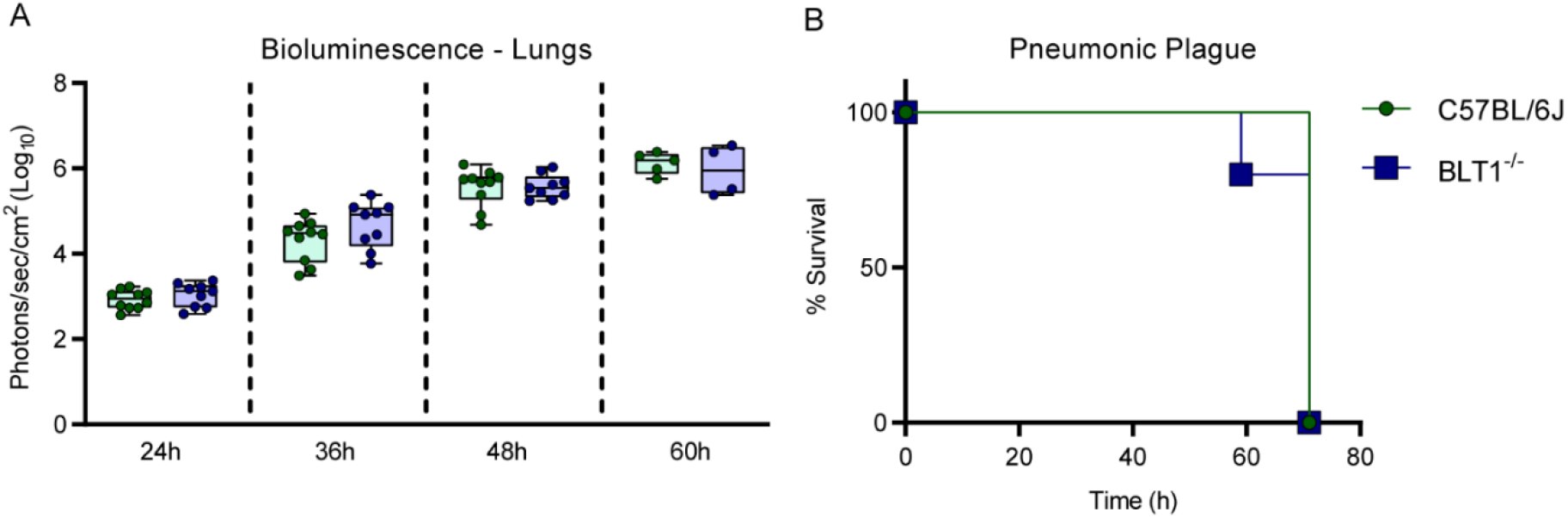
BLT1^-/-^ mice are not more susceptible to pneumonic plague than C57BL/6J mice. C57BL/6J and BLT1^-/-^ mice (n=10) were infected intranasally with 10x the LD_50_ of a bioluminescent strain of *Y. pestis* (*Y. pestis* CO92 LUX_P*cysZK*_) and monitored for proliferation by optical imagining and the development of moribund disease. (A) Bacterial proliferation in the lungs as a funciton of bioluminescence. (B) Survival curves of infected mice.

### Neutrophils do not synthesize LTB_4_ in response to *Y. pestis*

LTB_4_ is primarily produced by leukocytes such as mast cells, neutrophils, and macrophages (33). During pneumonic plague, *Y. pestis* initially interacts with alveolar macrophages, but by 12 h post-infection, the bacteria interact primarily with neutrophils (19). Moreover, we previously demonstrated that *Y. pestis* inhibits LTB_4_ synthesis by human neutrophils (48). Therefore, we sought to determine if murine neutrophils produce LTB_4_ in response to interactions with *Y. pestis*. Bone marrow-derived neutrophils from C57Bl/6J mice were infected with different gram-negative bacteria, and the synthesis of LTB_4_ was measured by ELISA. When stimulated with *E. coli, S. enterica* Typhimurium, or a strain of *K. pneumoniae* unable to synthesize its capsule, LTB_4_ synthesis was significantly induced within 1 h of infection (Fig. 3A; p≤ 0.0001). However, infection with *Y. pestis* did not elicit LTB_4_ synthesis, even when the MOI was increased to 100 bacteria per neutrophil (Fig. 3B). Similar phenotypes were observed during infection of human peripheral blood neutrophils, recapitulating our previous findings (Fig. 3C-D and (48)). Cell permeability and cytotoxicity were measured after infection to determine if the lack of LTB_4_ synthesis was due to *Y. pestis*-induced cell death. No significant cell permeability or cytotoxicity increases were observed during *Y. pestis* infections at an MOI of 20 (Fig. S1A-B). While slightly elevated permeability was observed at an MOI of 100 (Fig. S1C; 9% vs. 28%), overall cytotoxicity was lower in *Y. pestis* infected neutrophils than uninfected neutrophils (Fig. S1D; 4% vs. 12%). Similarly, *Y. pestis* did not induce elevated permeability or cytotoxicity in human neutrophils (Fig. S1E-F). These data demonstrate that neutrophils do not synthesize LTB_4_ in response to *Y. pestis*, and this phenotype is unique to this pathogen but not due to *Y. pestis* induced cell death.

**Fig 3.**
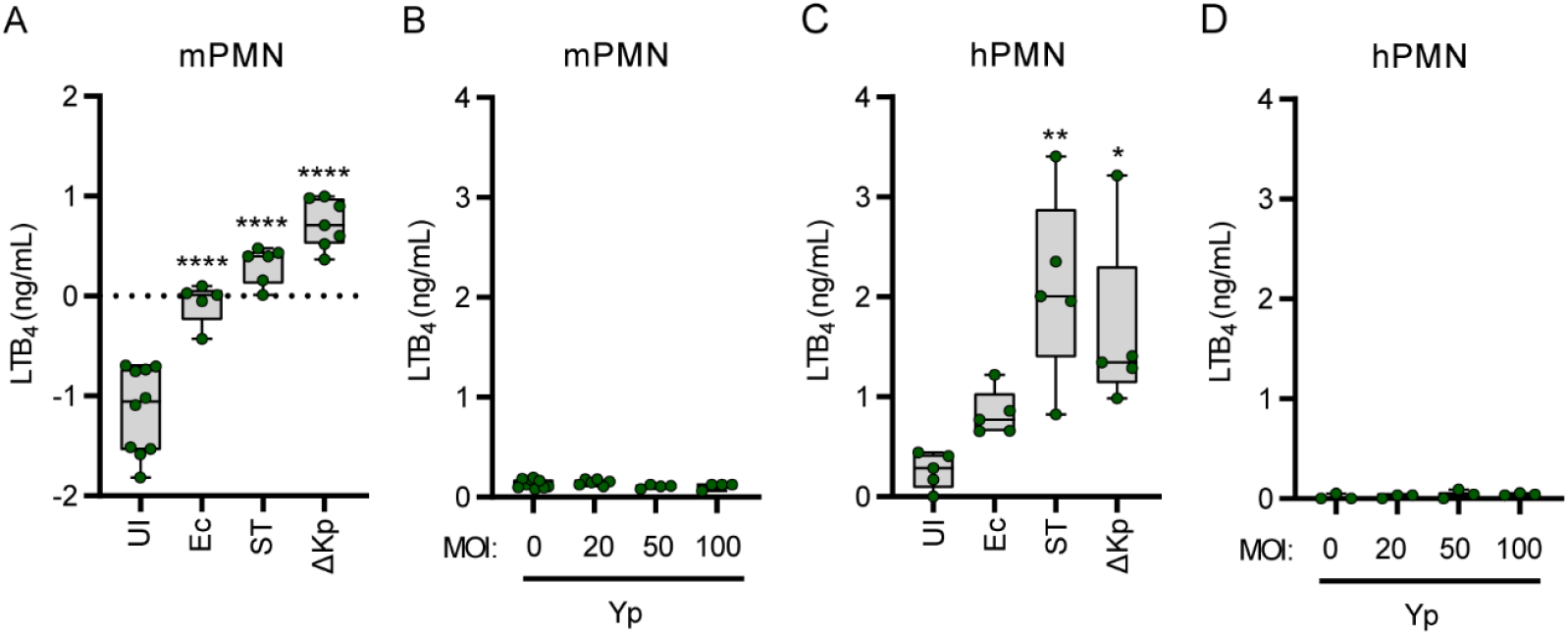
Neutrophils do not synthesize LTB_4_ in response to *Y. pestis*. (A-B) Murine or (C-D) human neutrophils were infected with *E. coli* DH5α (Ec), *S. enterica* Typhimurium LT2 (ST), or *K. pneumoniae manC* (ΔKp) at an MOI of 20, or with *Y. pestis* KIM1001 at increasing MOIs. LTB_4_ was measured from supernatants 1h post infection by ELISA. Each symbol represents independent biological replicates. UI=Uninfected. One-way ANOVA with Dunnett’s *post hoc* test. * = p≤0.05, ** = p≤0.01, **** = p≤0.0001.

### *Y. pestis* induces LTB_4_ synthesis in the absence of the Yop effectors

Seven Yop effector proteins are secreted into neutrophils via the T3SS (21-24), and we have previously shown Yop effector-mediated inhibition of LTB_4_ synthesis in human neutrophils at an MOI of 100 (48). However, Yop inhibition of LTB_4_ synthesis by murine neutrophils has not been previously shown and we wanted to independently confirm that the Yop effectors are sufficient to inhibit LTB_4_ synthesis at a lower MOI. Toward this goal, human and murine neutrophils were infected at an MOI of 20 with a *Y. pestis* mutant strain that expresses the T3SS but lacks all seven Yop effectors (*Y. pestis* T3E)(49). In contrast to *Y. pestis* infected cells, we observed a significant increase in LTB_4_ synthesis in response to the *Y. pestis* T3E strain, indicating that the Yop effectors are inhibiting synthesis (Fig 4A-B; p≤0.0001). To further determine which Yop effectors are required to inhibit LTB_4_ synthesis, neutrophils were infected with *Y. pestis* strains that expressed only one Yop effector (49). LTB_4_ synthesis was significantly decreased if *Y. pestis* expressed YpkA, YopE, YopH, or YopJ, and an intermediate phenotype was observed during infection with a strain expressing YopT (Fig. 4C). To demonstrate further that the Yop effectors were able to inhibit LTB_4_ synthesis actively, neutrophils were simultaneously infected with *Y. pestis* and the *Y. pestis* T3E mutant or with *Y. pestis* and a *K. pneumoniae* capsule mutant. Impressively, *Y. pestis* effectively abrogated LTB_4_ synthesis by neutrophils from both species stimulated by either *Y. pestis* T3E or *K. pneumoniae* (Fig. 4D-G). Together these data confirm that *Y. pestis* is not simply evading immune recognition but is actively inhibiting LTB_4_ synthesis via the activity of multiple Yop effectors.

**Fig 4.**
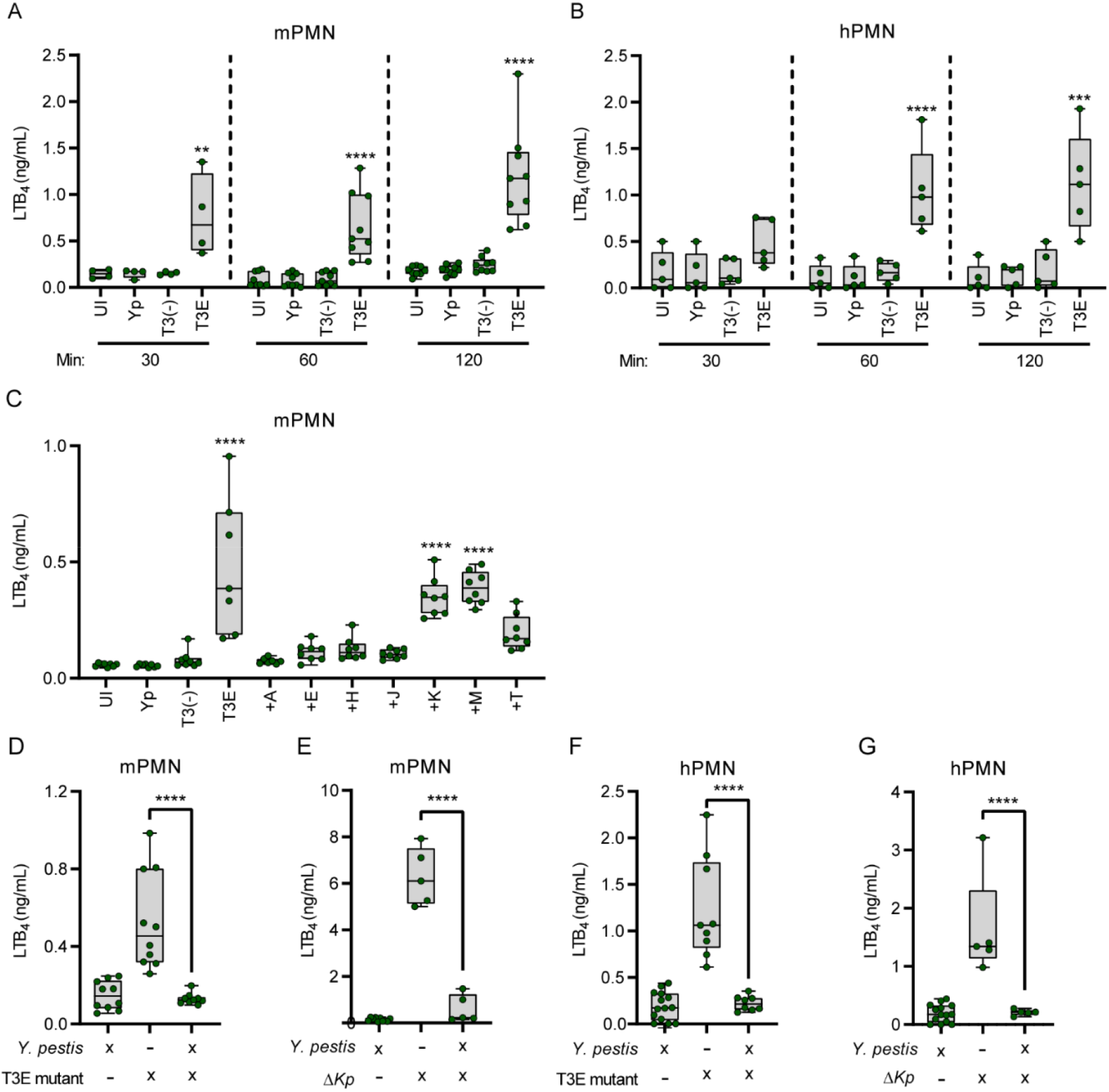
*Y. pestis* induces LTB_4_ synthesis in the absence of the Yop effectors. (A) Murine or (B) human neutrophils were infected with *Y. pestis* KIM1001 (Yp) or mutants that either lacked the Yop effector proteins (T3E) or the Yop effector proteins and the T3SS [T3(-)] (MOI of 20). LTB_4_ was measured from supernatants by ELISA at the indicated time points. (C) Murine neutrophils were infected with Yp, T3E, T3(-), or *Y. pestis* KIM1001 strains expressing only one Yop effector: +A = YpkA; +E = YopE; +H = YopH; +J = YopJ; +K = YopK; +M = YopM; or +T = YopT (MOI of 20). (D-E) Murine or (F-G) human neutrophils were co-infected with (D, F) Yp and T3E mutant or (E, G) with Yp and *K. pneumoniae manC* (ΔKp) (MOI of 20). LTB_4_ was measured from supernatants 1h post infection by ELISA. Each symbol represents independent biological replicates. UI=Uninfected. One-way ANOVA with Dunnett’s *post hoc* test. * = p≤0.05, ** = p≤0.01, *** = p≤0.001, **** = p≤0.0001.

### Neutrophils synthesize LTB_4_ in response to the *Y. pestis* T3SS in the absence of the Yop effectors

The T3SS is a pathogen-associated molecular pattern (PAMP) that is recognized by innate immune cells (7, 50, 51). To ascertain the role of T3SS in LTB_4_ synthesis by neutrophils during interactions with the *Y. pestis* T3E strain, we infected human and murine neutrophils with a *Y. pestis* strain lacking the pCD1 plasmid encoding the Ysc T3SS [*Y. pestis* T3^(-)^]. We did not observe any increase in LTB_4_ synthesis by neutrophils during interactions with *Y. pestis* T3^(-)^ compared to *Y. pestis*, even after 2 h of infection (Fig 4 A-B). Importantly, infection with the *Y. pestis* T3^(-)^ strain did not result in increased neutrophil cell permeability or cytotoxicity (S1 Fig.). To independently test that the T3SS is required to induce LTB_4_ synthesis, *Y. pestis* was grown under conditions that alter the expression of the T3SS prior to infection of neutrophils (5, 8, 16). Measuring expression of the LcrV protein as a proxy for overall T3SS expression confirmed decreased T3SS expression in cultures grown at 26°C compared to 37°C (Fig. 5A-B). As predicted by our pCD1 mutant data, LTB_4_ synthesis was not observed in *Y. pestis* strains grown at 26°C (Fig. 5C). Together, these data indicate that neutrophils recognize the *Y. pestis* T3SS as a PAMP that leads to the induction of LTB_4_ synthesis, but only in the absence of the Yop effectors.

**Fig 5.**
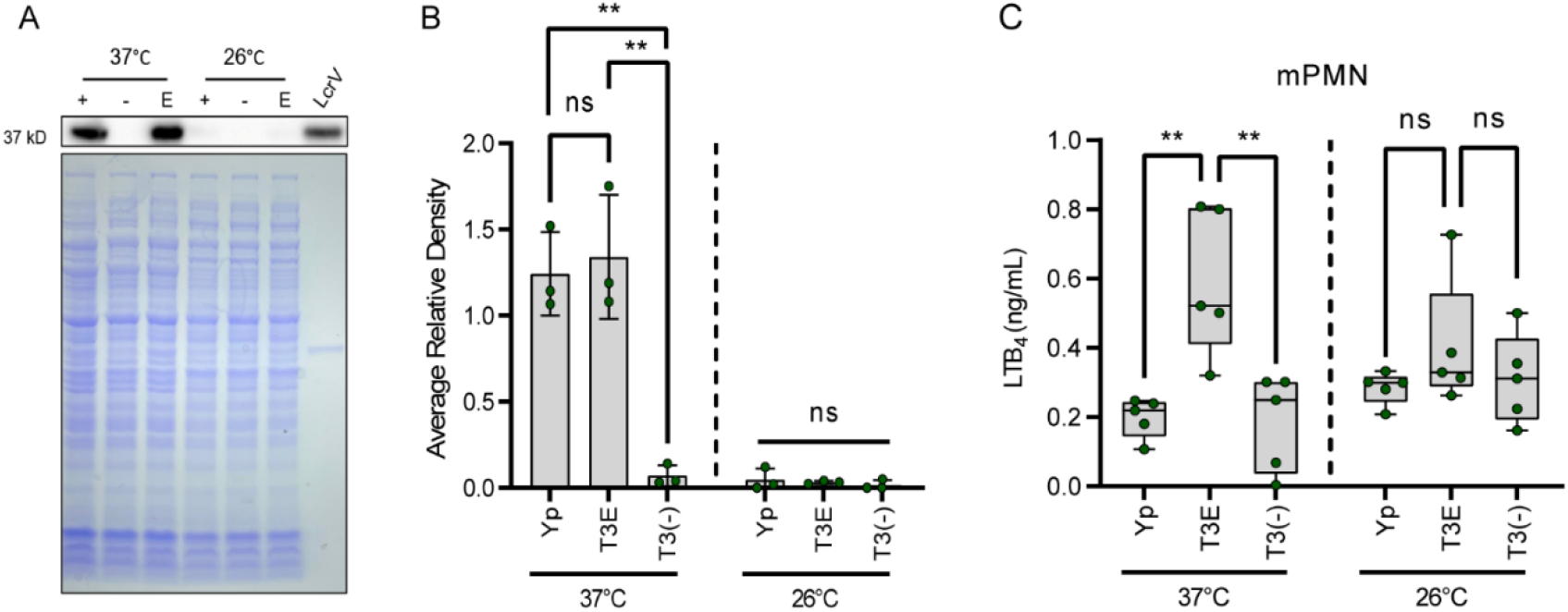
Neutrophils synthesize LTB_4_ in response to the *Y. pestis* T3SS in the absence of the Yop effectors. (A) Representative Western blot and coomassie images of *Y. pestis* KIM1001 lysates harvested from cultures grown at 37°C or 26 °C. (+) = Yp, (-) = T3(-); E = T3E; LcrV = 0.2 ug recombinant LcrV protein. (B) Average relative densities of LcrV from western blot images. (C) LTB_4_ measurement from supernatants of murine neutrophils infected for 1 h with *Y. pestis* KIM1001 (Yp) or mutants that either lacked the Yop effector proteins (T3E) or the Yop effector proteins and the T3SS [T3(-)] grown at 37°C or 26 °C (MOI of 20). Each symbol represents biological replicates. UI=Uninfected. One-way ANOVA with Tukey’s *post hoc* test. ns = not significant, ** = p≤0.01.

### *Y. pestis* inhibition of LTB_4_ synthesis is conserved during interactions with other leukocytes

In addition to neutrophils, two other lung resident leukocytes that can produce LTB_4_ are mast cells and, to a lesser degree, macrophages (31). To determine if *Y. pestis* inhibits LTB_4_ synthesis by these two cell types, bone marrow-derived mast cells and macrophages were isolated from C57Bl/6J mice and infected with *Y. pestis, Y. pestis* T3E, or *Y. pestis* T3^(-)^. As in neutrophils, we observed no synthesis of LTB_4_ by mast cells, even after 2 h of interacting with *Y. pestis* (Fig. 6A). However, LTB_4_ synthesis was significantly elevated in the absence of the Yop proteins (Fig. 6A, T3E; p≤0.01), reaching levels similar to that of mast cells stimulated with crystalline silica, a potent inducer of LTB_4_ synthesis in mast cells (52, 53). LTB_4_ synthesis by mast cells was similarly dependent on the presence of the T3SS, as the *Y. pestis* T3^(-)^ strain did not induce LTB_4_ synthesis (Fig. 6A). For macrophages, previous reports indicate that macrophage polarization influences the ability of macrophages to produce LTB_4_, with M1-polarized macrophages better able to synthesize LTB_4_ in response to bacterial ligands than M2-polarized cells (54). Therefore, we measured LTB_4_ synthesis of both M1- and M2-polarized macrophages. Again, we observed no significant synthesis of LTB_4_ by either macrophage population during interactions with *Y. pestis*, even after 4 h post-infection (Fig. 6A-B). However, significant synthesis of LTB_4_ was observed in M1-polarized macrophages in response to the *Y. pestis* T3E strain, which was dependent on the presence of the T3SS (Fig. 6B; p≤ 0.0001). As suggested by previous reports (55), we did not observe LTB_4_ synthesis by M2-polarized macrophages during interactions with any of the *Y. pestis* strains tested (Fig. 6C). Together, these data indicate that, as neutrophils, mast cells, and M1-polarized macrophages can quickly synthesize LTB_4_ in response to the *Y. pestis* T3SS, but the activity of the Yop effectors inhibits this response.

**Fig 6.**
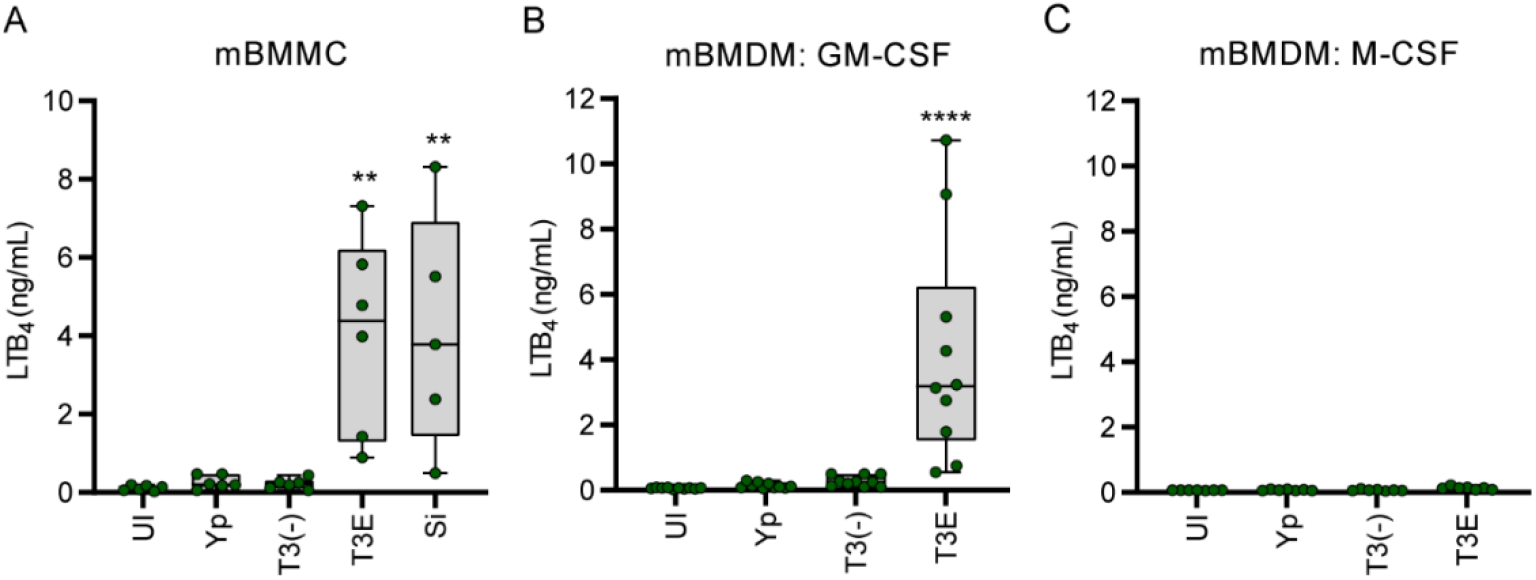
Lack of LTB_4_ response to *Y. pestis* is conserved in other leukocytes. (A)) Murine BMMCs were infected with *Y. pestis* KIM1001 (Yp), mutants that either lacked the Yop effector proteins (T3E), the Yop effector proteins and the T3SS [T3(-)], or treated with silica crystals (Si). LTB_4_ was measured from supernatants by ELISA at 2 h post infection. Murine BMDMs differentiated towards (B) M1 or (C) M2 phenotypes were infected with Yp, T3E, or T3(-). LTB_4_ was measured from supernatants by ELISA at 4 h post infection (MOI of 20). Each symbol represents independent biological replicates. UI=Uninfected. One-way ANOVA with Dunnett’s *post hoc* test. ** = p≤0.01, **** = p≤0.0001.

### Synthesis of PGE_2_ by neutrophils and macrophages, but not mast cells, is inhibited by *Y*. pestis

Unlike LTB_4_, the cyclooxygenase pathway appears to be induced during pneumonic plague (Fig. 1), suggesting that *Y. pestis* is unable to inhibit prostaglandin synthesis by leukocytes. Therefore, using PGE_2_ as a representative prostaglandin, we next examined the ability of murine neutrophils, macrophages, and mast cells to produce prostaglandins in response to *Y. pestis*. Like LTB_4_, neutrophils, and M1-polarized macrophages produce PGE_2_ in response to the T3SS, but synthesis is inhibited by secretion of the Yop effectors (Fig. 7A-B; p≤0.0001). However, mast cells appeared to produce equivalent amounts of PGE_2_ in response to all three strains of *Y. pestis*, indicating that *Y. pestis* is not able to inhibit PGE_2_ synthesis in mast cells (Fig. 7C; p≤0.05). These data suggest that signals leading to cyclooxygenase activity in mast cells differ from those in other leukocytes and that mast cells may be a primary source of PGE_2_ and other prostaglandins in response to *Y. pestis* infection of the lungs.

**Fig 7.**
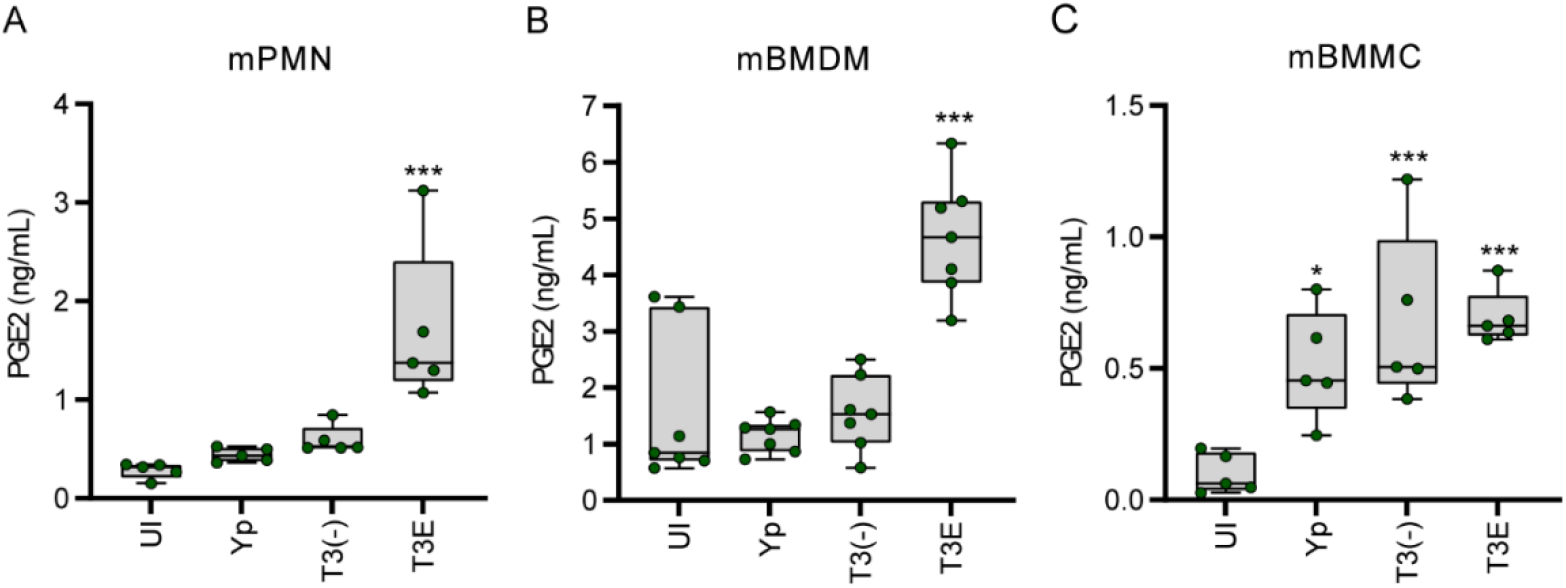
*Y. pestis* inhibits synthesis of PGE_2_ in neutrophils and macrophages but not mast cells. (A) Murine neutrophils, (B) M1-differentiated BMDMs, or (C) BMMCs were infected with *Y. pestis* KIM1001 (Yp) or mutants that either lacked the Yop effector proteins (T3E) or the Yop effector proteins and the T3SS [T3(-)]. PGE_2_ was measured from supernatants by ELISA at 1 h, 4 h, and 2 h post-infection, respectively. Each symbol represents independent biological replicates. UI=Uninfected. One-way ANOVA with Dunnett’s *post hoc* test. * = p≤0.05, *** = p≤0.001.

## Discussion

A hallmark manifestation of plague is the absence of inflammation during the early stages of infection, which is critical to *Y. pestis* virulence (10, 17, 26, 51). While *Y. pestis* has been shown to actively dampen the host immune response, there is a gap in our understanding of the role of lipid mediators of inflammation during plague. This study sought to define the host inflammatory lipid mediator response during pneumonic plague and expands our current understanding of how *Y. pestis* manipulates the immune system. During the earliest stages of infection, the host appears unable to initiate a timely LTB_4_ response. Because LTB_4_ is a potent chemoattractant crucial for rapid inflammation (29, 30, 56), this delay in LTB_4_ synthesis during plague likely has a significant impact on the ability of the host to mount a robust inflammatory response needed to inhibit *Y. pestis* colonization. First, in the absence of LTB_4_, sentinel leukocytes will not undergo autocrine signaling via LTB_4_-BLT1. Because LTB_4_-BLT1 engagement activates antimicrobial programs in leukocytes (29, 30, 45, 57-59), the absence of autocrine signaling diminishes the ability of sentinel leukocytes directly interacting with *Y. pestis* to mount an effective antimicrobial response to kill the bacteria. LTB_4_ synthesis is also regulated by BLT1 signaling, and autocrine signaling is required to amplify the production of LTB_4_ needed to rapidly recruit additional tissue-resident immune cells to the site of infection (29, 30, 58, 60, 61). Therefore, the normal feed-forward amplification of LTB_4_ synthesis, which is key for a rapid response to a bacterial infection, will also be inhibited by *Y. pestis*. Second, because LTB_4_ is required for neutrophil swarming (60, 62, 63), *Y. pestis* will also inhibit this key inflammatory mechanism (64). Neutrophil swarming is required to contain bacteria at initial sites of infection (65, 66). Thus, while individual neutrophils may migrate towards sites of *Y. pestis* infection, effective neutrophil swarming of large populations of neutrophils will be diminished. Finally, LTB_4_ is a diffusible molecule that can induce the inflammatory cascade in bystander cells (30, 67). Thus, while *Y. pestis* can inhibit cytokine and chemokine expression by cells with which it directly interacts (11, 15), inhibition of LTB_4_ synthesis likely also delays subsequent release of molecules by cells that do not directly interact with the bacteria. Together with the bacterium’s other immune evasion mechanisms, inhibition of LTB_4_ synthesis is likely another significant contributor to the generation of the non-inflammatory environment associated with the early stages of pneumonic plague (10, 11, 15). Incorporating these new LTB_4_ data with published findings from other laboratories (10, 11, 15), we have updated our working model of *Y. pestis* inhibition of inflammation during pneumonic plague (Fig. 8).

**Fig 8.**
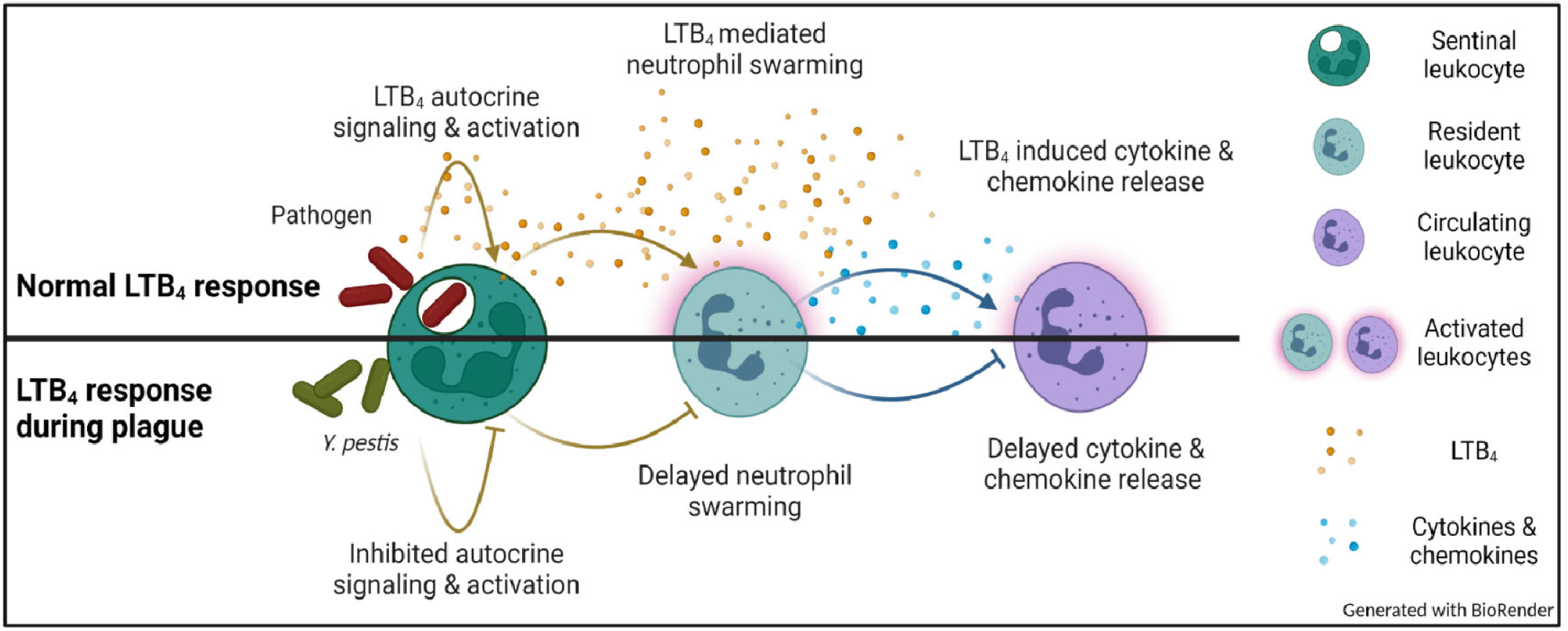
Working model for inhibition of the inflammatory cascade during plague.

These data also revealed that the T3SS translocon apparatus triggers LTB_4_ synthesis by leukocytes. Because our previous work with human samples indicated that neutrophils synthesize LTB_4_ in response to *Y. pestis* in the absence of the T3SS (48), we were initially surprised that we did not observe LTB_4_ synthesis by murine neutrophils to the *Y. pestis* T3^(-)^ strain. However, when we infected human neutrophils with lower MOIs, we observed that they also did not synthesize LTB_4_ in the absence of the T3SS (Fig. 4B, S2 Fig.). Under these infection conditions, neutrophils from both species only produced LTB_4_ in response to *Y. pestis* expressing the T3SS but none of the Yop effectors. These data support that the T3SS is a PAMP produced by *Y. pestis* that is not only recognized by macrophages (68) but also by neutrophils, which to our knowledge represents the first example of the *Y. pestis* T3SS serving as a PAMP in neutrophils. In macrophages, components of the T3SS are recognized by members of the nod-like receptor (NLR) family, leading to inflammasome activation (69-72), suggesting that inflammasome activation by the T3SS may trigger not only IL-1β/IL-18 secretion and pyroptosis but also LTB_4_ synthesis. However, whether inflammasome activation is required for the *Y. pestis* T3SS-mediated LTB_4_ synthesis remains unclear, as LTB_4_ synthesis is not always dependent on inflammasome activation (52, 73, 74). Interestingly, infection of neutrophils with a strain of *Y. pestis* that only expresses YopM, an effector that specifically inhibits the caspase-1 inflammasome (75), did not inhibit LTB_4_ synthesis (Fig. 4C and (48)), suggesting that LTB_4_ synthesis in response to the T3SS is not dependent on caspase-1 in neutrophils. Future studies using neutrophils from mice defective in specific NLRs and caspases will allow us to definitively determine if inflammasome activation is required for LTB_4_ synthesis in neutrophils in response to the *Y. pestis* T3SS. Moreover, because the enzymes that lead to LTB_4_ synthesis are well defined (32, 76), we can use *Y. pestis* mutants expressing different Yop effectors to specifically define the molecular mechanisms leading to activation of these enzymes, providing a clearer understanding of the signaling pathway(s) triggering LTB_4_ synthesis in the context of T3SS recognition. The lack of LTB_4_ synthesis in response to the *Y. pestis* T3^(-)^ strain also differed from what we observed for other gram-negative bacteria we tested (Fig. 4D-G), indicating that *Y. pestis* may also mask other potential gram-negative PAMPS that would typically be recognized by neutrophils. These data argue that *Y. pestis* has evolved both active (via the Yop effectors) and passive mechanisms to evade immune recognition and induction of LTB_4_ synthesis. Finally, it is worth noting that unlike human neutrophils, murine neutrophils did not appear to synthesize LTB_4_ during infections with the T3(-) strain at high MOIs (S2 Fig.). Differences in neutrophil responses between the two species have been well documented (77-81) but these observations merit further investigation into LTB_4_ responses by human neutrophils using higher MOIs to determine if human neutrophils are able to recognize other PAMPs during *Y. pestis* infection.

Finally, while we focused primarily on LTB_4_ in this study, our global lipidomics approach also revealed synthesis profiles for a variety of other inflammatory lipids that merit future considerations. The rapid cyclooxygenase response raises questions about whether prostaglandins are protective or detrimental during pneumonic plague. Historically, prostaglandins were thought to promote inflammation, but these mediators appear more nuanced under closer scrutiny and can just as likely inhibit inflammation as well as participate in normal development physiology without eliciting inflammation (39, 40, 82). All prostaglandins we observed as being significantly elevated during the non-inflammatory stage of pneumonic plague (PGA_2_, PGD_2_, PGE_2_, and PGJ_2_) have been shown to inhibit inflammation in various models, especially as synthesis levels increase (39, 40, 83-85). More specifically, PGE_2_ was shown to inhibit NADPH oxidase activity during infection with *Klebsiella pneumoniae*, which suppressed bacterial killing (86). PGE_2_ has also been shown to directly counteract the proinflammatory activities of LTB_4_ (87, 88). The phagocytic index of LTB_4_-stimulated rat alveolar macrophages (AMs) is reduced when co-stimulated with PGE_2_ (88). Moreover, AMs treated with PGE_2_ show a 40% reduction in LTB_4_ synthesis when stimulated with an ionophore known to induce a strong LTB_4_ response (87). This inhibition of LTB_4_ by PGE_2_ is suspected to be via an increase in second messenger cAMP that activates protein kinase A (PKA), which has been shown to inhibit LTB_4_ synthesis (87, 89). Together, these data suggest that the elevated levels of prostaglandin synthesis observed during pneumonic plague may also contribute to the blunted LTB_4_ response we observed during pneumonic plague. Interestingly, our in vitro data indicate that *Y. pestis* also inhibits prostaglandin synthesis by macrophages and neutrophils but not mast cells, suggesting that mast cells may be the primary source of prostaglandins during pneumonic plague. Surprisingly, while mast cells are important sentinel leukocytes in the lung and dermis, their contributions during plague and responses to *Y. pestis* have not been previously explored. Our discovery that mast cells respond differently to *Y. pestis* than other leukocytes support that we need more studies into the role of these cells during plague.

In conclusion, we have defined the kinetics of the inflammatory lipid mediator response during pneumonic plague, which revealed a blunted LTB_4_ response during the early stages of infection. Furthermore, we have shown that *Y. pestis* actively manipulates lipid synthesis by leukocytes via the activity of Yop effectors to generate a beneficial inflammatory outcome to the pathogen. These discoveries warrant further research into the role of lipids, and subsequent manipulation of their synthesis by *Y. pestis*, to fully understand the molecular mechanisms *Y. pestis* has evolved to manipulate the mammalian immune response.

## Material and Methods

### Bacterial strains

Bacterial strains used in these studies are listed in S2 Table. For mouse infections, *Y. pestis* was grown at 26 °C for 6-8 h, diluted to an optical density (OD) (600 nm) of 0.05 in Bacto brain heart infusion (BHI) broth (BD Biosciences, Cat. No. 237500) with 2.5 mM CaCl_2_ and then grown at 37 °C with aeration for 16 to 18 h (90). For cell culture infections, *Y. pestis* was cultured with BHI broth for 15 to 18 h at 26°C in aeration. Cultures were then diluted 1:10 in fresh warmed BHI broth containing 20 mM MgCl_2_ and 20 mM Na-oxalate and cultured at 37°C for 3 h with aeration to induce expression of the T3SS. Bacterial concentrations were determined using a spectrophotometer and diluted to desired concentrations in 1 × Dubelco’s phosphate-buffered saline (DPBS) for mouse infections or fresh medium for *in vitro* studies. Concentrations of bacterial inoculums for mouse studies were confirmed by serial dilution and enumeration on agar plates.

### Mouse infections

All animal work was reviewed and approved by the University of Louisville IACUC prior to initiation of studies and performed twice to ensure reproducibility. Infected mice were monitored for the development of moribund disease symptoms twice daily and humanely euthanize when they met previously approved end point criteria. C67BL/6J or BLT1^-/-^ mice (91)(6-8 weeks) were anesthetized with ketamine/xylazine and administered 20 μL bacteria suspended in 1× DPBS to the left nare as previously described (47, 90). For lipidomic measurements, mice were humanely euthanized by CO_2_ asphyxiation at 6, 12, 24, 36, or 48 h and lungs were harvested. Lungs were transferred to a 2 mL tube pre-filled with 2.8 mm ceramic beads (VWR, Cat. No. 10158-612), flash frozen on dry ice, and stored at −80°C until preparation for lipidomic analysis. For BLT1^-/-^ studies, mice were infected with a strain of *Y. pestis* carrying a bioluminescent bioreporter to monitor bacterial proliferation and dissemination by optical imaging using an IVIS Spectrum In Vivo Imaging System (Caliper) as previously described (47).

### Lipid extraction and quantification by LC-MS

To prepare the samples for lipidomic analysis, first lungs were thawed with 1.5 mL of ice cold 1 X DPBS +HALT protease and phosphatase inhibitor cocktail for 3 minutes. Lungs were then homogenized with Bead Ruptor 4 (OMNI) at speed 5 (5 m/s) for 3 cycles of 30 seconds with 1-minute pauses in which the lungs were placed on ice. Tissue debris was then centrifuged for 10 min at 1,500 x g at 4°C. The supernatant (∽1.5 mL) was then transferred to a fresh eppendorf tube. From this, 250 μL of supernatant were combined with 750 μL of 100% methanol + 0.1% BHT (final concentration of 75%) and incubated at 4°C for 24 h to inactivate *Y. pestis* and extract lipids. After confirmation of successful inactivation of *Y. pestis*, lipids were extracted an quantified by the Wayne State University Lipidomics Facility as previously described (38). Briefly, samples were applied to conditioned C18 reverse phase cartridges, washed with water followed by hexane and dried under vacuum at the end of each wash. Cartridges were then eluted with 1 mL methanol containing 0.1% formic acid. The eluate was dried under a gentle stream of nitrogen. The residue was redissolved in 30 μL methanol that was diluted with 30 μl of 10 mM aqueous ammonium acetate and readied for LC–MS analysis. The extracted samples were analyzed for the fatty acyl lipidome using standardized methods as described earlier (92, 93).

### Cell isolation and cultivation

Use of human neutrophils was approved by the University of Louisville Institutional Review Board (IRB) guidelines (approval no. 96.0191). Human neutrophils were isolated from the peripheral blood of healthy, medication-free donors, as described previously (94). Neutrophil isolations yielded ≥95% purity and were used within 1 h of isolation. Murine neutrophils were isolated from bone marrow of 7–12-week-old mice using an Anti-Ly-6G MicroBeads kit (Miltenyi Biotec; Cat. No. 130-120-337) per the manufacturer’s instructions. Neutrophil isolations yielded ≥95% purity and were used within 1 h of isolation. Macrophages were differentiated from murine bone marrow in DMEM supplemented with 30% L929 conditioned media, 1 mM Na-pyruvate, and 10% FBS for 6 days. Macrophages were either polarized with 10 ng/mL of GM-CSF (Kingfisher Biotech; Cat. No. RP0407M) or with 10 ng/mL of M-CSF (Kingfisher Biotech; Cat. No. RP0462M) throughout the differentiation. The medium was replaced on days 1 and 3. Murine mast cells were isolated and differentiated from bone marrow as previously described (95). Briefly, isolated bone marrow cells were resuspended in BMMC culture medium [DMEM containing 10% FCS, penicillin (100 units/mL), streptomycin (100 mg/mL), 2 mmol/L L-glutamine, and 50 mmol/L β-mercaptoethanol] supplemented with recombinant mouse stem cell factor (SCF) (12.5 ng/mL; R&D Systems, Cat. No. 455-MC) and recombinant mouse IL-3 (10 ng/mL; R&D Systems, Cat. No. 403-ML). Cells were plated at a density of 1 × 10^6^ cells/mL in a T-75 cm^2^ flask. Nonadherent cells were transferred after 48 hours into fresh flasks without disturbing the adherent (fibroblast) cells. Mast cells were visible after 4 weeks of culture and propagated further or plated for experiments in DMEM without antibiotics.

### Leukocyte infections

Human neutrophils (1 × 10^6^) were resuspended in Kreb’s buffer (w/ Ca^2+^ & Mg) then adhered to 24-well plates that were coated with pooled human serum for 30 min prior to infection (wells were washed twice with 1 x DPBS prior to plating the cells). Murine neutrophils (1 × 10^6^) were resuspended in RPMI + 5% FBS then adhered to 24-well plates that were coated with FBS for 30 min prior to infection (wells were washed twice with 1 x DPBS prior to plating the cells). Neutrophils were infected at a multiplicity of infection (MOI) of 20, 50, or 100 and incubated for 1 h in a 37°C CO_2_ cell culture incubator. Co-infections were performed at a final MOI of 20 (10 for each strain). 1 h post-infection, supernatants were collected, centrifuged for 1 min at 6,000 x g’s, and supernatants devoid of cells were transferred to a fresh eppendorf tube. Samples were stored at −80°C until ELISA analysis. Macrophages (2 × 10^6^) were adhered to 24-well plates in DMEM + 10% FBS 1 day prior to infection. Macrophages were infected at an MOI of 20. At 4 h post infection, supernatants were collected, centrifuged for 1 min at 6,000 x g’s, and supernatants devoid of cells were transferred to a fresh eppendorf tube. Samples were stored at −80°C until ELISA analysis. Mast cells (2.5 × 10^5^) were adhered to 24-well plates in DMEM only for 1 h prior to infection. Mast cells were infected at an MOI of 20 or treated with crystalline silica (100 mg/cm^2^). At 2 h post infection supernatants were collected, centrifuged for 1 min at 6,000 x g’s, and supernatants devoid of cells were transferred to a fresh eppendorf tube. Samples were stored at −80°C until ELISA analysis.

### Measurement of LTB_4_ and PGE_2_ by enzyme-linked immunosorbent assay

Supernatants of neutrophils, macrophages, and mast cells were collected and measured for LTB_4_ or PGE_2_ by enzyme-linked immunosorbent assay (ELISA) per manufacturer’s instructions (Cayman Chemicals; Cat. No. 520111 and Cat. No. 514012, respectively).

### Cell viability assays

To determine leukocyte permeability, cells were incubated with 90% trypan blue for 5 min and trypan blue exclusion was measured using SD100 counting chambers (VWR; Cat. No. MSPP-CHT4SD100) and a cell counter (Nexcelom Cellometer Auto T4). To determine leukocyte cytotoxicity, lactate dehydrogenase (LDH) was measured from leukocyte supernatants using CytoTox 96 Non-Radioactive Cytotoxicity kit (Promega; Cat. No. g1780) per manufacturer’s instructions.

### Measurement of LcrV by western blot

Bacterial strains were cultured with BHI broth for 15 to 18 h at 26°C in aeration. Cultures were then diluted 1:10 in fresh warmed BHI broth containing 20 mM MgCl_2_ and 20 mM Na-oxalate and cultured at 37°C or 26°C for 3 h. 1 OD_600_ of bacterial pellets were collected and resuspended in SDS-PAGE loading buffer, boiled for 10 min, and 0.1 OD_600_ was separated on a 10% SDS-PAGE gel. As a positive control, 0.2 g of recombinant LcrV protein (BEI resources; Cat. No. NR-32875) was used. Samples were immunoblotted with polyclonal anti-LcrV antibody (BEI Resources; Cat. No. NR-31022) diluted to 1:4,000. Anti-goat IgG HRP secondary antibody was diluted to 1:5,000 (Bio-Techne; Cat. No. HAF017). Densitometry was performed using ImageJ software to compare LcrV bands between samples (96).

### Statistics

Human neutrophils were harvested from both male and female donors and infections were performed on different days. Murine experiments were performed on both male and female mice and were performed on different days.Where appropriate, one-way analysis of variance (ANOVA) with Dunnett’s or Tukey’s post-test, or T-test with Mann-Whitney’s *post*-test, as indicated in individual figure legends, were used for statistical analysis, and performed using Prism 8 (GraphPad). For the LC-MS data, a LIMMA-Moderated t-test was performed using a modified version of our previously published protocol using R packages (97-99). Briefly, raw data were transformed by taking logarithmic base 2 followed by quantile normalization. Missing values were then ascribed using a singular value decomposition method. Lipids missing > 40% of the values were excluded from subsequent analysis. Finally, differentially abundant lipids (p<0.05) were further filtered by fold-change (FC) criteria (1 < log_2_FC < 1) and multiple comparisons testing with a false discovery rate.

## Acknowledgements

The authors would like to acknowledge Dr. Jon Goguen for generously sharing the *Yersinia pestis* KIM1001 strains.

## Supporting Information

**S1 Fig.**
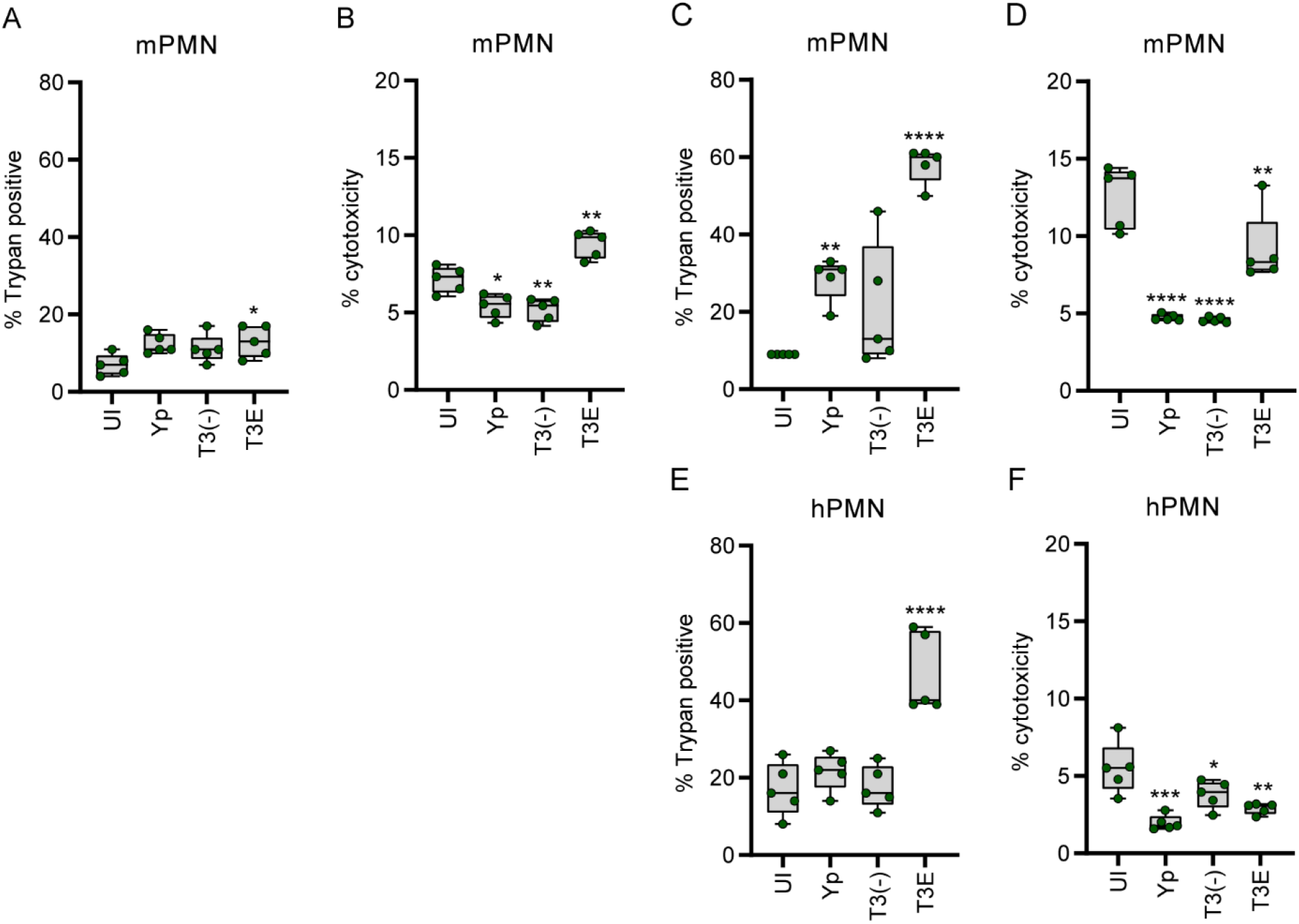
Absence of LTB_4_ response to *Y. pestis* is not due to cell death. (A-D) Murine or (E-F) human neutrophils (∽95% purity) were infected with *Y. pestis* KIM1001 at (A-B) an MOI of 20 or (C-F) an MOI of 100 for 1 h. (A, C, E) Cell permeability as a function of trypan exclusion. (B, D, F) Cytotoxicity as a function of LDH release. UI=Uninfected. (A-F) One-way ANOVA with Dunnett’s *post hoc* test. * = p≤0.05, ** = p≤0.01.

**S2 Fig.**
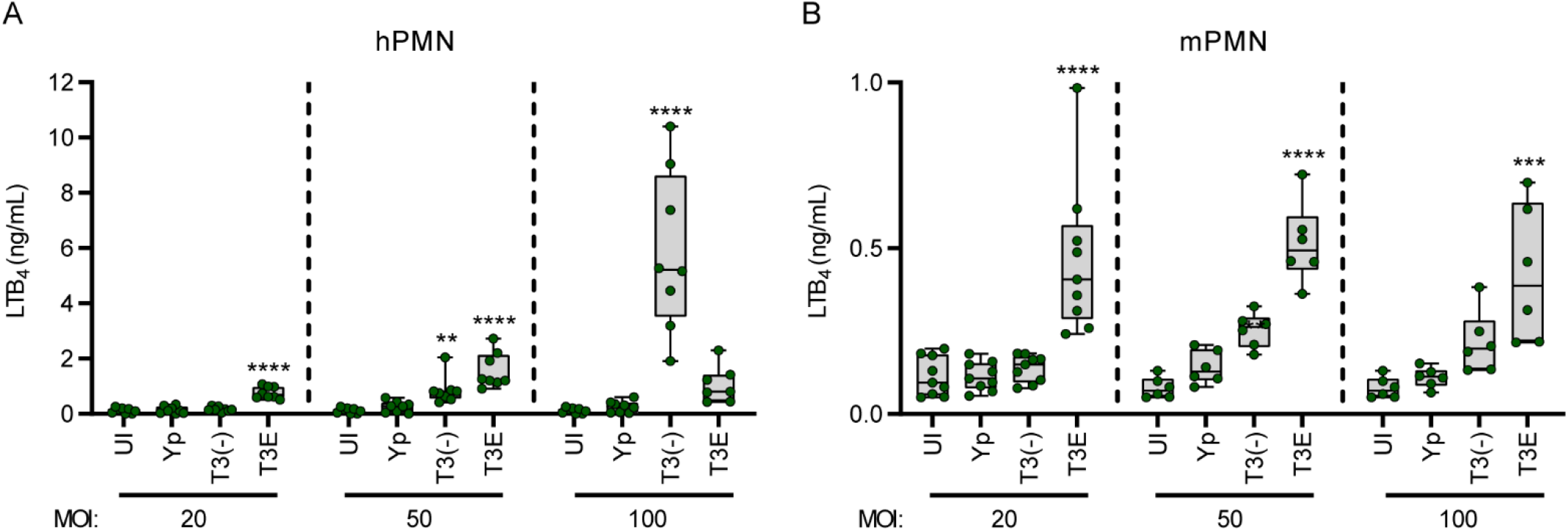
Differential recognition of T3^(-)^ *Y. pestis* between human and mice neutrophils. (A) Human or (B) murine neutrophils (∽95% purity) were infected with *Y. pestis* KIM1001 or mutants that either lacked effector proteins (T3E) or lacked effector proteins and the T3SS (T3(-)) at increasing multiplicities of infections (MOIs). LTB_4_ was measured from supernatants by ELISA 1 h post infection. Each symbol represents independent biological replicates. UI=Uninfected. (A-B) One-way ANOVA with Dunnett’s *post hoc* test. ** = p≤0.01, *** = p≤0.001, **** = p≤0.001.

**S3 Fig.**
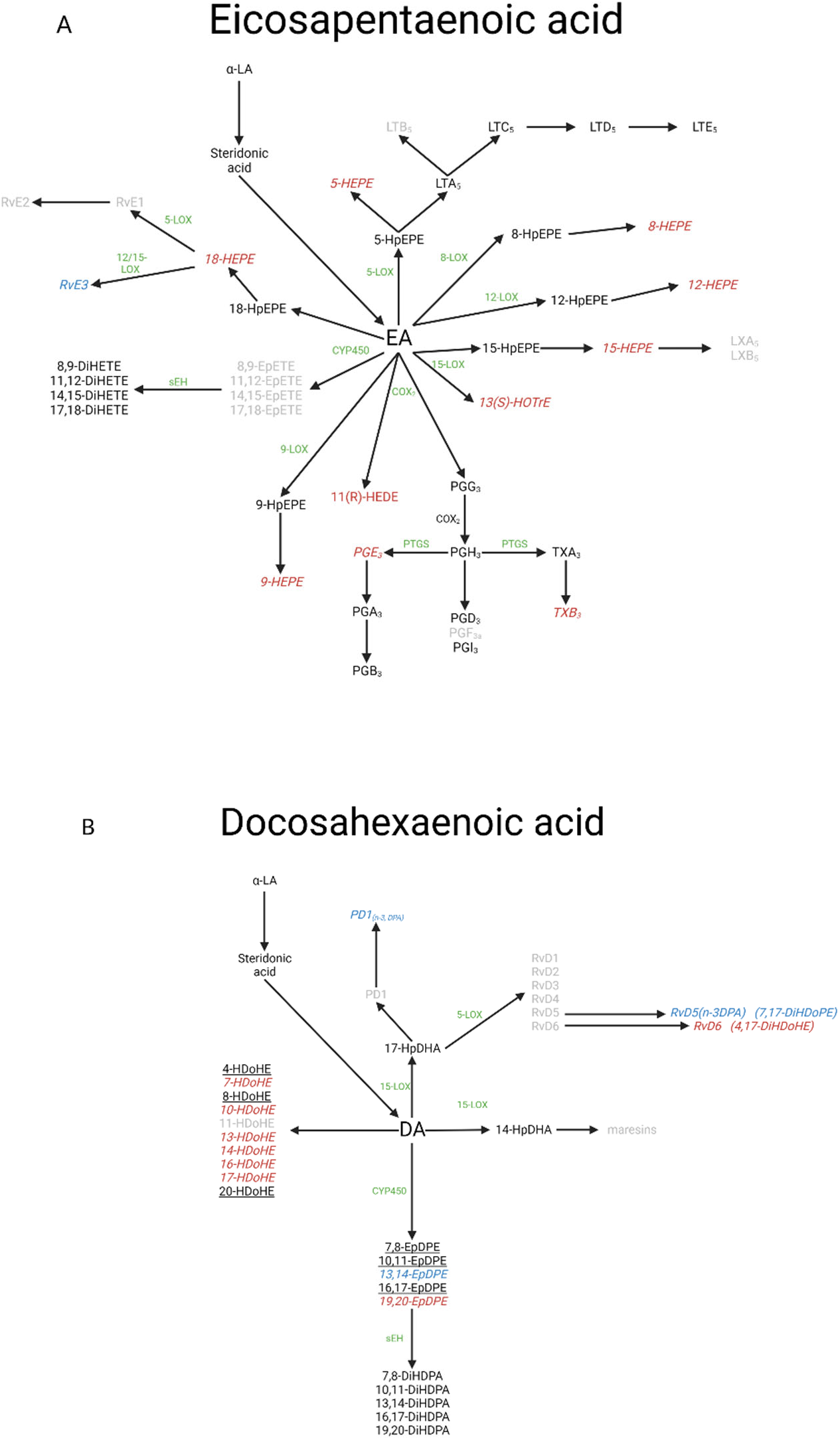
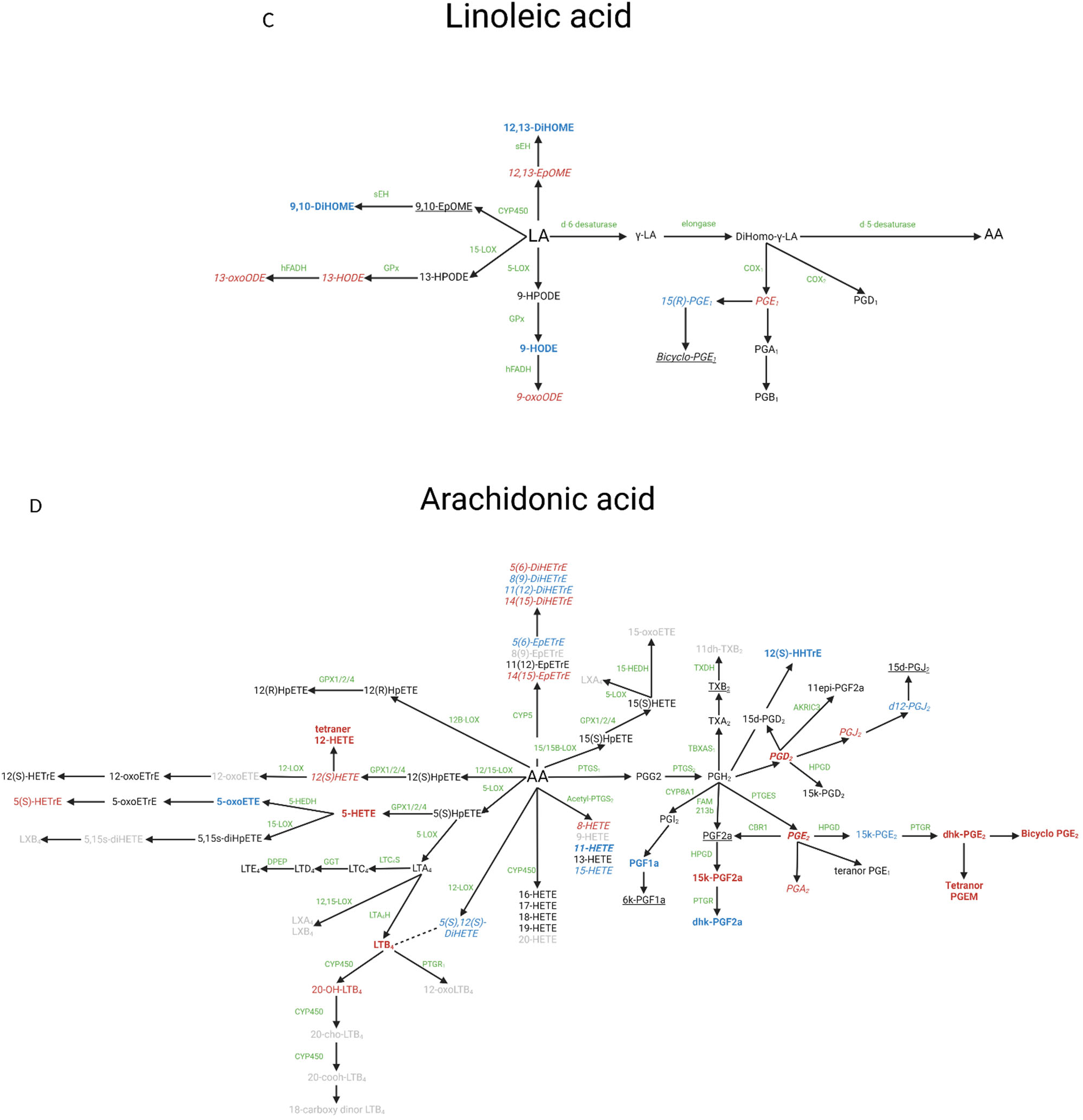
Synthesis pathways of eicosanoids. (A) Eicosapentaenoic acid, (B) docosahexaenoic acid, (C) linoleic acid, and (D) arachidonic acid pathways and the products measured in LC-MS. Black – Not screened; Red – significant increase compared to uninfected in at least one time point; Blue – significant decrease compared to uninfected in at least one time point; Grey – below the limit of detection; Green – enzyme responsible for lipid conversion (no enzyme indicates a non-enzymatic conversion via redox); Underlined – no change; Dotted line – epimers. Of the significant hits: Bold-pro-inflammatory; Italicized-anti-inflammatory/pro-resolving.

**S1 Table. Changes in inflammatory lipids during first 48h of pneumonic plague**. C57Bl/6J mice were infected with 10X the LD_50_ of *Y. pestis* KIM5 and lungs were harvested at 6, 12, 24, 36, and 48 h post-infection (n=5). Total lipids were isolated from homogenized lungs and 143 lipids were quantified by LC-MS. Significant changes in lipid concentrations were observed in at least one time point for 63 lipids.

**S2 Table.**
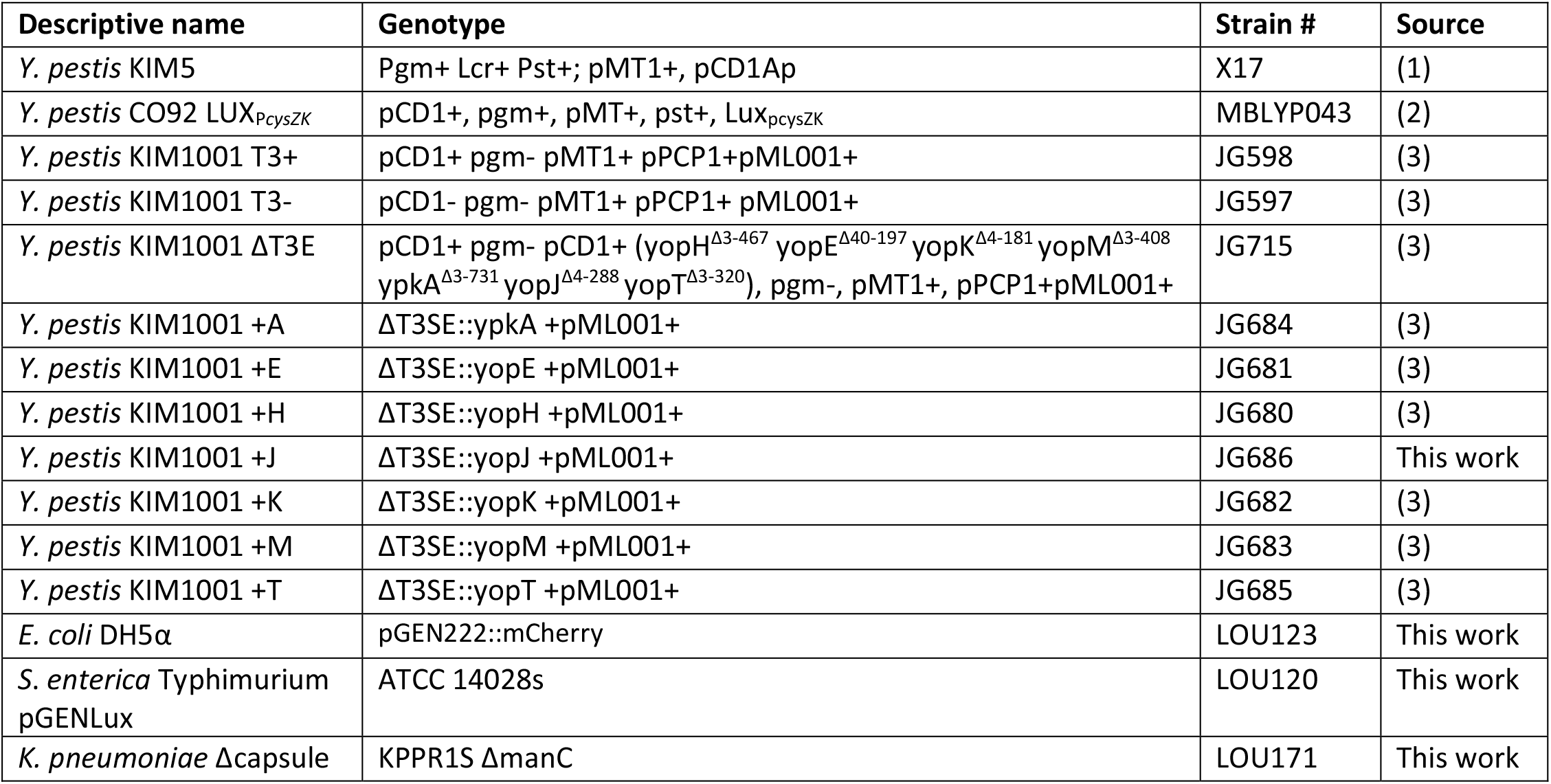
Bacterial Strains used in this study.

| Descriptive name | Genotype | Strain # | Source |
| --- | --- | --- | --- |
| *Y. pestis* KIM5 | Pgm+ Lcr+ Pst+; pMT1+, pCD1Ap | X17 | (1) |
| *Y. pestis* CO92 LUX<sub>pcysZK</sub> | pCD1+, pgm+, pMT+, pst+, Lux<sub>pcysZK</sub> | MBLYP043 | (2) |
| *Y. pestis* KIM1001 T3+ | pCD1+ pgm- pMT1+ pPCP1+pML001+ | JG598 | (3) |
| *Y. pestis* KIM1001 T3- | pCD1- pgm- pMT1+ pPCP1+ pML001+ | JG597 | (3) |
| *Y. pestis* KIM1001 ΔT3E | pCD1+ pgm- pCD1+ (yopH<sup>Δ3-467</sup> yopE<sup>Δ40-197</sup> yopK<sup>Δ4-181</sup> yopM<sup>Δ3-408</sup> ypkA<sup>Δ3-731</sup> yopJ<sup>Δ4-288</sup> yopT<sup>Δ3-320</sup>), pgm-, pMT1+, pPCP1+pML001+ | JG715 | (3) |
| *Y. pestis* KIM1001 +A | ΔT3SE::ypkA +pML001+ | JG684 | (3) |
| *Y. pestis* KIM1001 +E | ΔT3SE::yopE +pML001+ | JG681 | (3) |
| *Y. pestis* KIM1001 +H | ΔT3SE::yopH +pML001+ | JG680 | (3) |
| *Y. pestis* KIM1001 +J | ΔT3SE::yopJ +pML001+ | JG686 | This work |
| *Y. pestis* KIM1001 +K | ΔT3SE::yopK +pML001+ | JG682 | (3) |
| *Y. pestis* KIM1001 +M | ΔT3SE::yopM +pML001+ | JG683 | (3) |
| *Y. pestis* KIM1001 +T | ΔT3SE::yopT +pML001+ | JG685 | (3) |
| *E. coli* DH5α | pGEN222::mCherry | LOU123 | This work |
| *S. enterica* Typhimurium pGENLux | ATCC 14028s | LOU120 | This work |
| *K. pneumoniae* Δcapsule | KPPR1S ΔmanC | LOU171 | This work |

